# Avian haemosporidians (*Plasmodium* and *Haemoproteus*) status in selected bird groups (Old world Flycatchers, Warblers, Babblers, and Thrushes) of India and their phylogenetic relationships with other lineages of the world

**DOI:** 10.1101/2021.09.27.461904

**Authors:** Vipin, Ashutosh Singh, Rajnikant Dixit, Narinder Kumar Tripathi, Bhim Singh, Vinita Sharma, Chandra Prakash Sharma, Dhananjai Mohan, Sandeep Kumar Gupta

**Author notes:** **Address for correspondence**, Dr. S. K. Gupta, Scientist E, Wildlife Institute of India, Chandrabani, Dehradun, 248 001 (U.K.), India.

## Abstract

The avian haemosporidians (*Plasmodium* and *Haemoproteus*) are widely distributed and may affect the host populations from body damage at individual level to the extinction of a population. The knowledge about their status may help in future avifauna conservation plans. Hence, we examined the avian haemosporidians status, in selected bird groups (Old world Flycatchers, Warblers, Babblers, and Thrushes) of India, and their phylogenetic relationships with other known lineages of the world. We used the common genetic marker (Cytochrome *b* gene fragment of 479 bp) with information on the geographic distribution of parasite and host species available at MalAvi database. The prevalence of avian haemosporidians, from northern, eastern, and southern parts of India and phylogenetic genetic analysis of lineages was carried out to know the genetic relatedness among them at local and world level. The MCC tree revealed six Haemosporidian lineages in which one was common (H_MSP01) and five were unique (H_CYOPOL01, H_CHD01, H_CYORUB01, H_EUMTHA01, and P_GEOCIT01). The avian host richness Index was 2.0852. 9.9%, prevalence of *Haemosporidian* infection was found in 111 DNA samples belonging to 6 host species. The *Haemoproteus* prevalence was found to be 9.0 % across five host species (*Phylloscopus trochiloides, Cyornis poliogenys, C. hainanus dialilaemus, C. rubeculoides, Eumiyas thalassinus*) and *Plasmodium* prevalence was 0.9% in *Geokichla citrina*. Spatial phylogeny at global level showed H_MSP01 lineage, found in different host species in India, was genetically related to *H*. *pallidus* lineages (COLL2 and PFC1) in parts of Africa, Europe, North America, Malaysia, and Philippines. The *Plasmodium* lineage (P_GEOCIT01) was related to PADOM16 in Egypt with poor sequence similarity (93.89%). The statistical analysis suggested that the haemosporidian’s host species distribution range was directly and significantly associated with the altitude, minimum temperature, and relative humidity. H_MSP01 distribution was in accordance with *H. pallidus* having a wide geographic and host range.

## Introduction

*Plasmodium* and *Haemoproteus* (Apicomplexa) are common and widely distributed vector-borne blood parasites, which occur in different bird species (1, 2). These haemosporidians (*Plasmodium* and *Haemoproteus*) may affect the avian host species by damaging their tissues (3, 4) during the stages of their life cycle (5, 4), by decreasing the survival rate (4, 6, 7, 8), lowering their reproductive success (4, 9, 10) and affecting their body conditions negatively (4, 11, 12). The haemosporidian infections have been reported to impart more severe effects on the avian populations in terms of reduction, extermination and even extinction of their populations (4, 13, 14). Hence, considering the mega diversity status of India where more than thirteen percent of worlds avifauna is found (http://bnhsenvis.nic.in/files/Bird_Diversity_Popular_Lecture.pdf), it becomes much important to study the status of avian haemosporidians to ensure the proper conservation strategies of different host species in future.

Though many molecular studies have been carried out on the avian haemosporidians in the world (4), but there is record of only few such studies from India in which one is confined to a single bird species (15) and the second is particular to certain mountain ranges (16). While studying the avian haemosporidians at the genetic level, the lack of common genetic markers and information on the geographic distribution of parasite and host samples makes it exceedingly difficult to compare the data (17). Further, the uncommon lineage naming and use of a common length of a standard genetic marker by researchers make it very difficult to carry out a fair comparison of data at the end (18, 19). Therefore, many studies, these days, are making use of the MalAvi database, which is available at http://mbio-serv4.mbioekol.lu.se/avianmalaria/index.html for download (20, 21, 22). This database is developed by the partial fragments of cytochrome b gene primer pairs for *Plasmodium*, *Haemoproteus* and *Leucocytozoon* parasites (17). It contains information on these avian blood parasites and their hosts’ geographic locations (23). Thus, removing the above-mentioned problems.

In the present study we describe the status of Haemosporidians (*Palsmodium* and *Haemoproteus*) in selected bird groups (Old world Flycatchers, Warblers, Babblers, and Thrushes) of India. The Old-world Flycatchers studied in the current study belonged to the families of Muscicapidae, Timaliidae, Stenostirdae, Phylloscopidae and Turdidae. The Muscicapidae and Turdidae families have a worldwide distribution, are strongly migratory, and winter in South-east Asia. The Warblers belong to the family Phylloscopidae are distributed in Eurasia, Wallacea, Africa, and Alaska and winter in Southeastern Asia. The Babblers belong to the family Timaliidae and are mostly distributed in the Indian Sub-continent and in Southeast Asia. So, with the help of the information retrieved from the MalAvi database, we have tried to locate the genetically related Haemosporidian lineages with current samples and their hosts through spatial phylogenetics. The present study considered the vast distributional range of hosts, from the extreme north to south and east of India. The status of only *Plasmodium* and *Haemoproteus* was studied in selected bird groups in India as these are the two most common genera of parasites found in avian blood (20). The sampling regions represent a large range of altitudinal variation, relative humidity, and temperature which may be affecting the host and parasite distribution. We also attempted to explain any correlation that exists among these parasites.

## Materials and Methods

### Sample collection

DNA from blood and tissue samples of birds (n=111), belonging to the group of Old world Flycatchers (2 Families and 14 species), Warblers (1 Family and 1 species), Babblers (1 Family and 2 species), and Thrushes (1 Family and 1 species), collected from the western Himalayas, Northeastern Hill states and Eastern Ghats of India between the period of 2014 to 2018 for a different project at Wildlife Institute of India, Dehradun were used for the current study. The distribution and sampling details about each sample are provided in Fig. 1 and Table S1.

**Figure 1.**
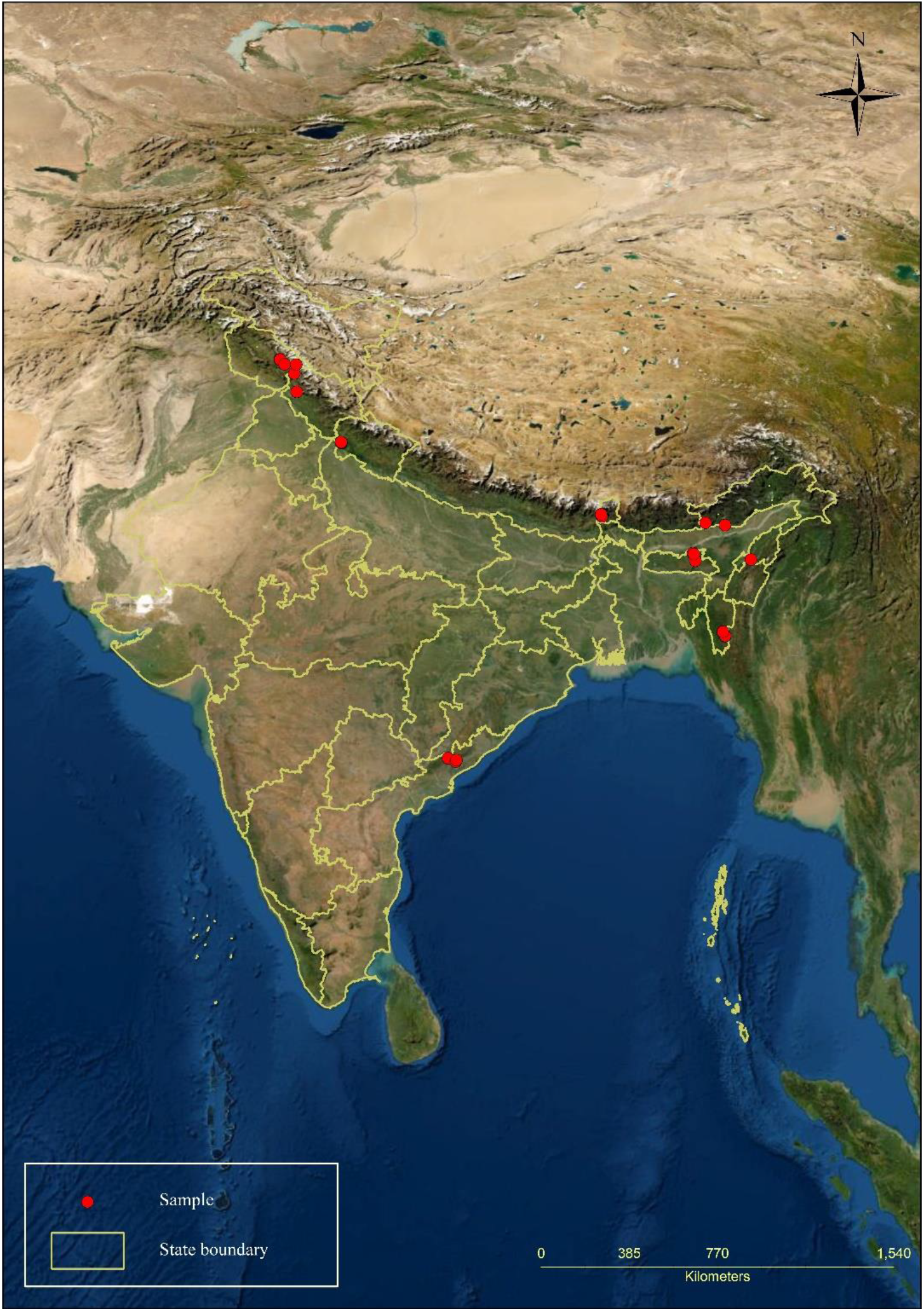
The distribution details of collected bird samples.

### DNA extraction and host species identification

The genomic DNA was extracted using DNeasy blood and tissue kit (Qiagen, Hilden, Germany) as described by Singh et al (24), and species identification was done as described previously by Singh et al (25). All DNA extraction steps were carried out in a sterile and dedicated laboratory to avoid any contamination. We also included one negative extraction control to check any cross-contamination in the kit reagents.

### Identification of Plasmodium and Haemoproteus parasite lineages

To identify the lineages of *Plasmodium* and *Haemoproteus*, the 479 bp region of the Cytochrome *b* gene was amplified using the primers HAEMF and HAEMR2 (23). These primers were chosen because the region between them has been proven to show the almost haplotypes that could be present in the entire cytochrome *b* gene (26). The *Plasmodium* DNA was used as positive control while performing all PCR reactions. The PCR amplification was carried out in a 10-μl reaction volume containing 8 μl of Taq PCR Master Mix (Qiagen, Hilden, Germany), 10 μM of each primer (1 μl), and 1 μl of the genomic DNA. The cycling conditions were initial denaturation at 95 °C for 5 min, followed by 35 cycles of denaturation at 95 °C for 40 s, annealing at 53 °C for 45 s, and extension at 72 °C for 45 s, with a final extension for 10 min at 72 °C in a Veriti Thermal Cycler (Applied Biosystems, Foster City, CA). We also amplified the *Plasmodium* genus-specific fragment of cytochrome b gene (596 bp) with primer pair L1 and L2 (27) in all DNA samples to cross check the PCR amplification failure by HAEMF and HAEMR2 primers. The reaction volumes and PCR conditions for L1 and L2 were as described by the Tan et al (27). The amplified PCR products were visualized on 2% agarose gel. Before sequencing, the PCR products were treated using exonuclease I and shrimp alkaline phosphatase enzyme to remove the residual primer and dNTPs. The sequencing was carried out in both the directions using BigDye RR kit 3.1 (Applied Biosystems, Foster City, CA) in a 3500xL Genetic Analyzer (Applied Biosystems, Foster City, CA). The generated sequences were cleaned manually on Sequencing Software v5.1 (Applied Biosystems, Foster City, CA). The sequences were aligned using BioEdit Sequence Alignment Editor Software v7.2.5 (28). The PCR and DNA sequencing were carried out at Wildlife Institute of India, Dehradun. The sequences were BLAST searched on the MalAvi database for avian haemosporidian parasites (29). A parasite sequence was defined as a unique lineage if a single nucleotide difference was found while comparing it with the sequences available at MalAvi database (23, 30, 31)

### Phylogenetic analysis

The Bayesian evolutionary analysis of *Plasmodium* and *Haemoproteus* lineages was done using BEAST2 v2.6.2 (32) to know the level of genetic relatedness among them. The host phylogenetic tree was constructed using the generated cytochrome b sequences. The phylogenetic relationships of the lineages, from the current study, at the global level were analyzed by obtaining the cytochrome *b* sequences of a total of 2862 *Plasmodium* and *Haemoproteus* lineages, of which 1287 lineages were of *Plasmodium* and 1575 lineages were of *Haemoproteus* from the MalAvi database for avian haemosporidian parasites, as of 24-05-2020, in FASTA format (Table S2A). Three lineages of Leucocytozoon (L_CLAHYE01, L_CLAHYE02, L_CLAHYE03) were used as outgroups. The HKY substitution model was selected with Gamma Category Count 4. The empirical frequencies were selected for getting a good fit to the data. The strict molecular clock model was selected for the data. The Yule Model of speciation was selected for the tree prior as it has been considered more appropriate when using different species sequences. The priors for rest parameters were set as default. The MCMC chain length was set to 6 million for this data set. The trace log and tree log frequencies were set to 1000 and the screen log frequencies were set to 10000. Assessment of adequate sampling and other of BEAST output parameters was done with TRACER (33). To obtain the phylogenetic tree, the time trees produced by the BEAST were summarized using TreeAnnotator program v1.7.2. (34). The TreeAnnotator burns the first 10% of the total 6000 trees generated by BEAST and the maximum clade credibility (MCC) tree with the highest posterior probability of all nodes was generated. The annotated tree was visualized on the FigTree program v1.4.4 (35).

### Spatial phylogenetic analysis of Plasmodium and Haemoproteus lineages

The Bayesian phylogeographic analysis of *Plasmodium* and *Haemoproteus* lineages of this study was carried out to know their phylogenetic relationships with other known lineages of these two genera at the global level using Spherical Biogeography package with BEAST v2.6.2 (36). For this, the cytochrome *b* sequences (n=54) of all lineages (excluding not having geographical coordinates information) of *Plasmodium* (n=1) and *Haemoproteus* (n=53) (Table S3), showing maximum similarity percentage with current study lineages, were used from the MalAvi database (obtained on 24-05-2020) for comparison (Table S4). The Geo Sphere package having a phylogeographic model was installed in the BEAST. The tip dates were set and the HKY substitution model was selected with empirical frequencies. A strict clock model was set. The Coalescent with Constant Population model was chosen for the tree prior. The information for latitude and longitude of each lineage was entered in the Spherical Geography model. The clock rate was set to 2e-5 for the data set the partition to speed up the convergence. For the geography partition, we selected a relaxed clock with a log-normal distribution. The MCMC chain length was set to 1 million. The screen log and file log were set to 10000 and 1000, respectively. The tree annotation was done as mentioned above, except the first 10% of the total 1000 trees were burned. The maximum clade credibility file was KML rendered using SpreaD3 (Spatial Phylogenetic Reconstruction of Evolutionary Dynamics) program v.0.9.7.1rc (37) for spatial map construction on Google Earth Pro. The DNA sequence similarity percentage was calculated through Multiple Sequence Alignment using MUSCLE (38).

### Species richness, the correlation between physical parameters with host and parasite distribution, if any

Margalef Richness Index for Avian host species was calculated using the formula = (S - 1) / Log (n), where S = total number of species and n = total number of individuals in the sample. Further statistical analysis was carried out by taking the host species Groupwise i.e., Old world Flycatcher, Warblers, and Thrushes. The values of elevation for each sample were recorded by GPS as described by Kumar et al (39). Other physical parameters viz. relative humidity, minimum, and maximum temperature for each collection site and date were obtained from NASA’s meteorological data sets available from Data Access Viewer (https://power.larc.nasa.gov/data-access-viewer/) (Table S1). The elevation range (minimum and maximum) for each avian host found infected with parasites was obtained from the IUCN Red List of Threatened Species (https://www.iucnredlist.org) (Table S1). The linear correlation, regression, and Karl Pearson’s Correlation were carried out for physical parameters (relative humidity, minimum, and maximum temperature and elevation range) of the study area from where avian host samples were found infected with blood parasites using SPSS statistics software version 19 (40).

## Results

### Parasite lineages and their phylogeny analysis at world level

Of the 111 host DNA samples, the cytochrome *b* gene’s sequencing was done for 17 parasite positive samples out of which only 11 sample’s (LM1, LM2, LM3, LM4, LM6, LM10, LM16, AS86, AS89, AS91, and AS102) sequences belonging to six species (*Phylloscopus Trochiloides, Cyornis poliogenys, Geokichla citrina, Cyornis hainanus dialilaemus, Cyornis rubeculoides* and *Eumyias thalassinus*) were usable (Table 1). The cytochrome *b* gene fragment of 596 base pairs was also amplified with L1 and L2 primers for the above 17 samples. The ten sequences representing five lineages were matched with *Haemoproteus* and one sequence with *Plasmodium* and thus a total of six lineages were found (Table 1). The naming of these six lineages was done as suggested by Bensch et al (17), which were as follows H_MSP01, H_CYOPOL01, H_CHD01, H_CYORUB01, H_EUMTHA01, and P_GEOCIT01. The maximum clade credibility tree of six lineages has been shown in Fig. 2. The genetic unrelatedness of *Haemoproteus* and *Plasmodium* lineages of this study are supported by high values of tree nodes posterior probabilities (0.696-0.999), (Fig. 2). Out of 479 base pairs of cytochrome *b* gene the consensus sequences of 230 base pairs for the 6 lineages could be drawn after the alignment. The aligned consensus sequences of these 6 lineages were used to show their relationships with all available lineages of *Haemoproteus* and *Plasmodium* on the MalAvi database. The phylogenetic analysis at the global level was carried out by making a consensus sequence of 230 base pairs of cytochrome *b* sequences of a total of 2686 *Plasmodium* and *Haemoproteus* lineages, of which 1230 lineages were of *Plasmodium* and 1456 lineages were of *Haemoproteus* (Table S2B).

**Table 1.**
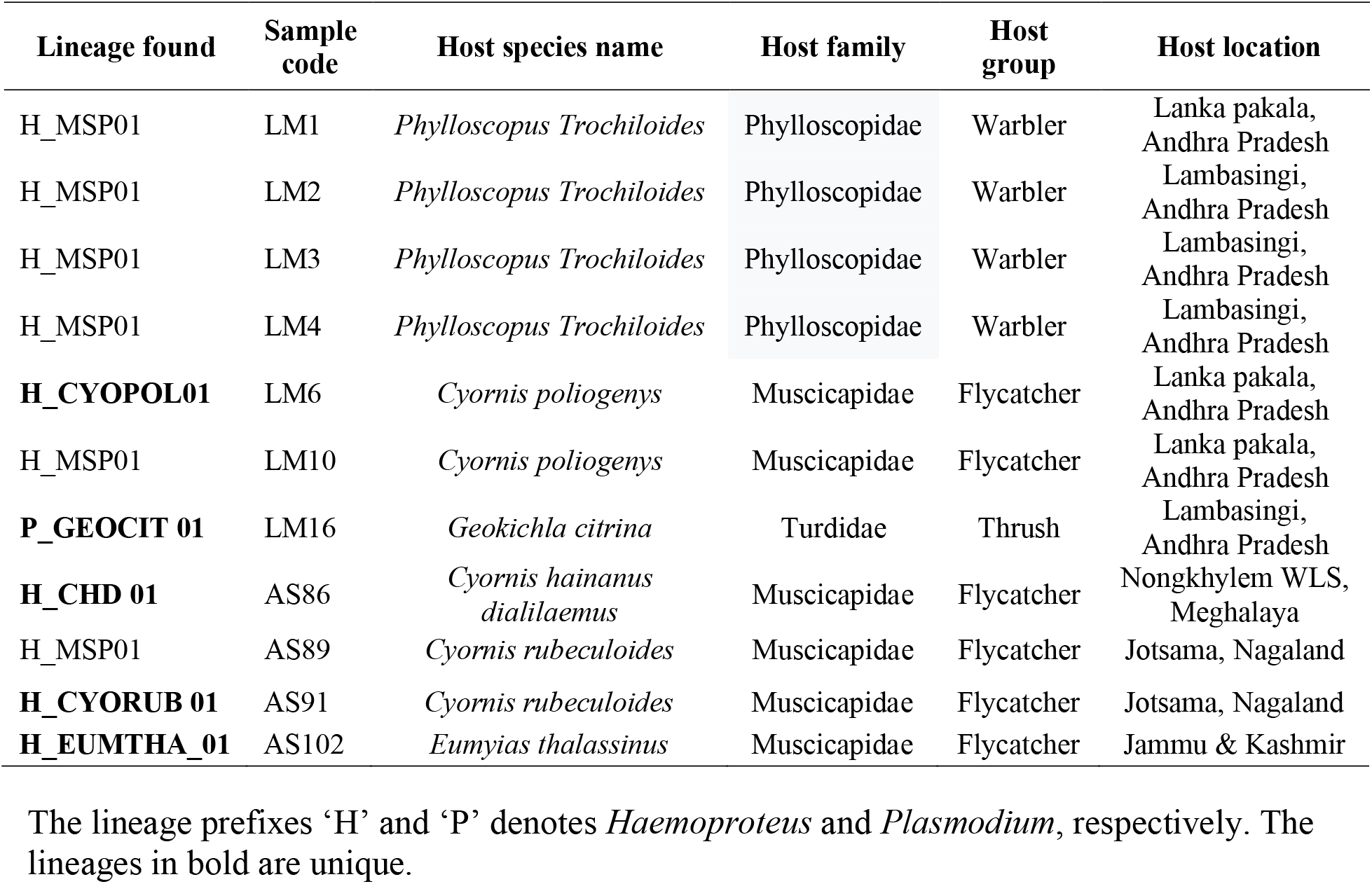
Lineages of *Plasmodium* and *Haemoproteus* found in different host species in the present study.

**Figure 2.**
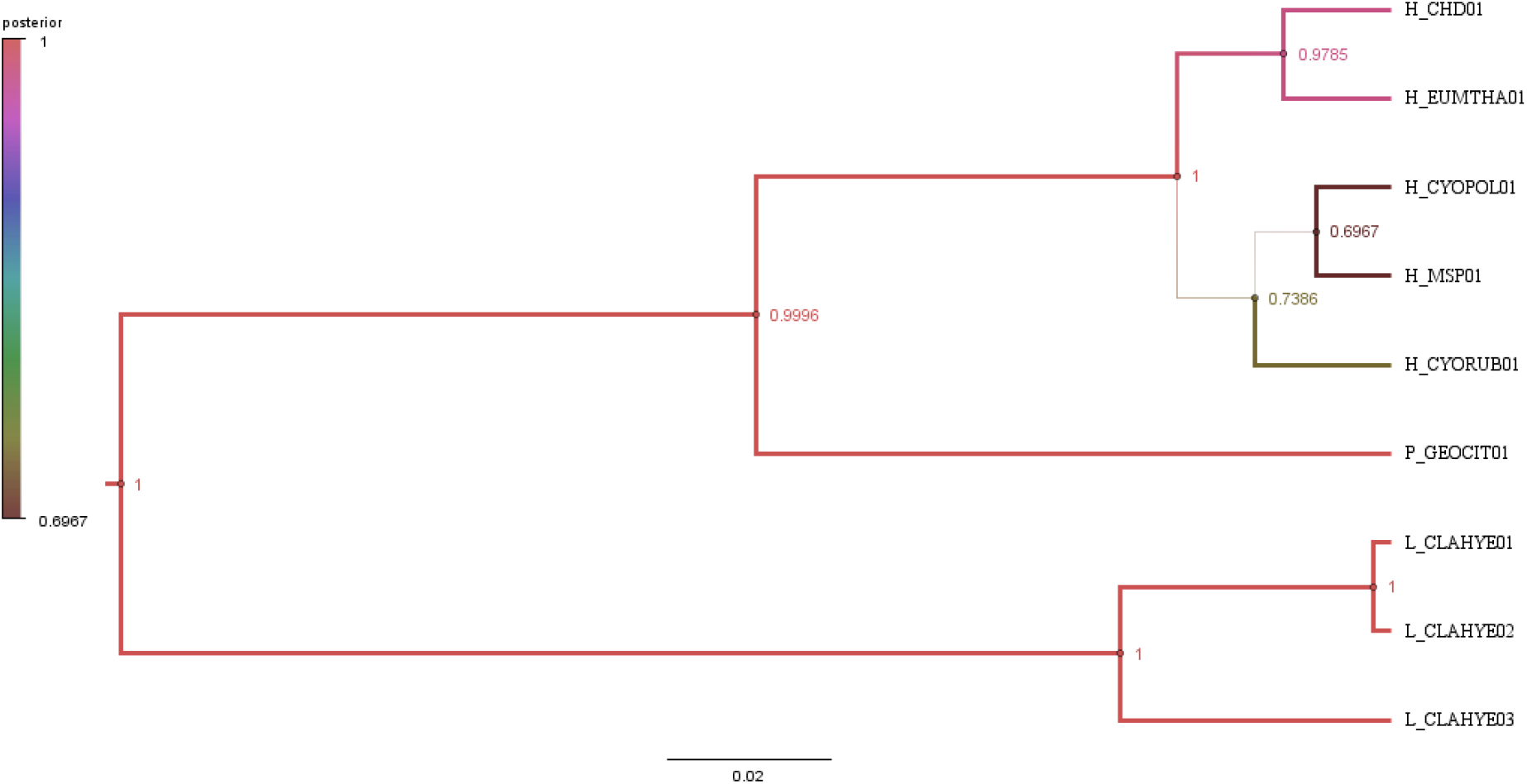
The maximum clade credibility tree showing six lineages. The node labels represent the posterior probabilities. Node values, shapes, tree branches, legend colour and line width are coloured and given, respectively, as per posterior probability values.

In world phylogenetic tree the clades, other than representing the samples of current study, were collapsed to make the tree small and for adequate representation. The phylogenetic analysis of five *Haemoproteus* lineages revealed that the clade having MSP01 lineage clustered with clades containing COLL2 and PFC1 lineages, which belong to *Haemoproteus pallidus* (Posterior probability 0.056) (Fig. 3). Further, MSP01 shared 100% sequence similarity with COLL2 and PFC1 (Table S4). The clades containing the lineages CYOPOL01, EUMTHA01, and CYORUB01 were same having the COLL2 and PFC1 lineages. The CHD01 lineage clustered with lineage TROERY02 in a different clade (Fig. 3), however, their sequence similarity was very low (97.38%) (Table S4). Lineage GEOCIT clustered PADOM16 in the *Plasmodium* clade and shared the maximum sequence similarity with it at a global level i.e., 93.89% (Fig. 3 and Table S4). Excluding MSP01, the rest five lineages matched 93% to 99% with the MalAvi database, whose details have been given in Table S4. So, all lineages in this study except MSP01 were unique. The short sequence of 230 base pairs do not hinder, here, in defining a new lineage as enough variations within this fragment, different from all known lineages available at MalAvi database, were found.

**Figure 3.**
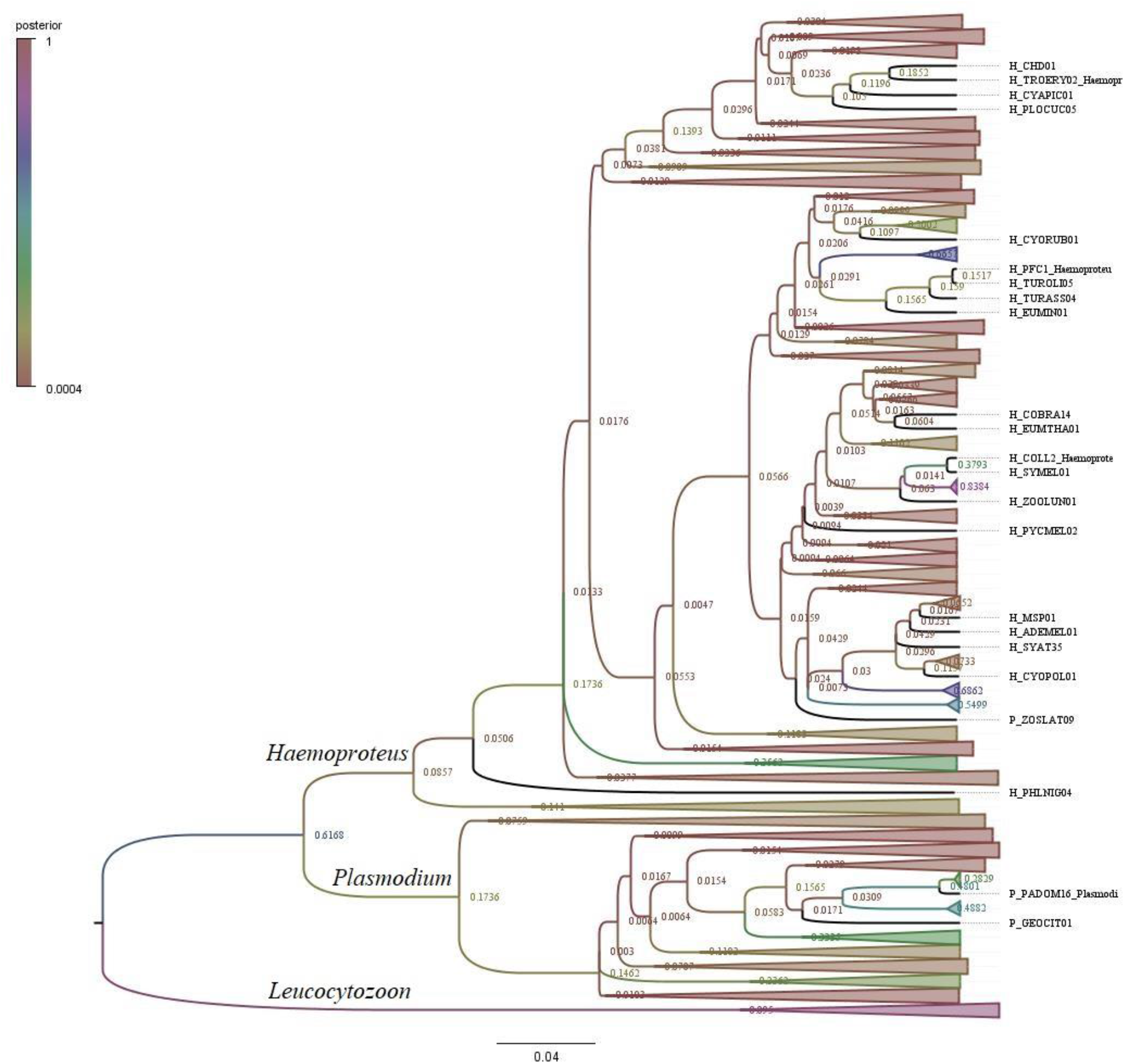
The maximum clade credibility tree showing World phylogeny of all *Plasmodium* and *Haemoproteus* lineages, including from this study. The node values, legend, clade and the colour of all these parameters are set as per the posterior probability values.

### Parasite prevalence and host-parasite association

Of the 111 DNA samples belonging to the eighteen host species, haemosporidian infection was found in 11 hosts belonging to the six species, which showed a 9.9% prevalence. *Haemoproteus* prevalence was found to be 9.0% across five host species (*Phylloscopus trochiloides, Cyornis poliogenys, C. hainanus dialilaemus, C. rubeculoides, Eumiyas thalassinus*) (Table 1). *Plasmodium* prevalence was 0.9% in one species of bird (*Geokichla citrina*). The *Haemoproteus* haplotype MSP01 was more common in warbler’s species *Phylloscopus trochiloides* (N=4) than in two species of Old-World Flycatchers *C. poliogenys* and *C. rubeculoides* (N=2). The *Haemoproteus* lineage CYOPOL01, CHD01, CYORUB01, and EUMTHA01 were unique to *C. poliogenys, C. hainanus dialilaemus, C. rubeculoides, and Eumiyas thalassinus, respectively*. The *Plasmodium* lineage GEOCIT01 was unique to the species *Geokichla citrina*.

### Correlation between physical parameters with host and parasite distribution

The Margalef Avian host richness Index was calculated to be 2.0852. The correlation between relative humidity and altitudinal height was found to be positive (R^2^ = 0.77) (Table 2). The regression model predicted that the altitude and minimum temperature values showed R^2^ = 0.77 and significant (*p* < 0.003) (Fig. 4). The correlation coefficients among different physical parameters are shown in Table 3. A significantly high negative correlation (87.8%) was observed between altitude and minimum temperature (Table 3), suggesting that this parameter is directly related to the host’s distribution range. Using Pearson’s correlation analysis test, different physical parameters were associated with the occurrence of host species in different locations sampled. The 3D scatter plot summarizes all the above parameters collectively (Fig. 5).

**Table 2.**
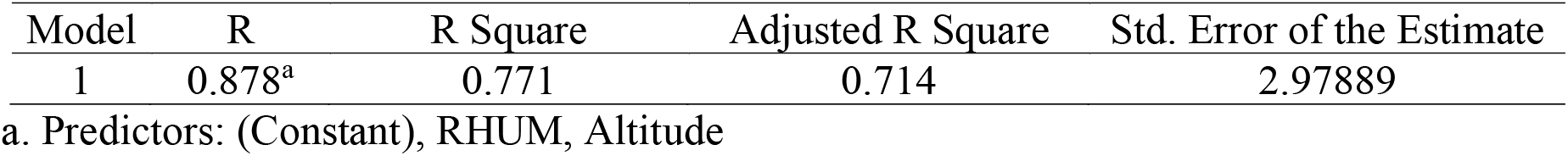
Model Summary showing simple corelation and total variation (humidity and altitude).

| Model | R | R Square | Adjusted R Square | Std. Error of the Estimate |
| --- | --- | --- | --- | --- |
| 1 | 0.878<sup>a</sup> | 0.771 | 0.714 | 2.97889 |

a. Predictors: (Constant), RHUM, Altitude

**Figure 4.**
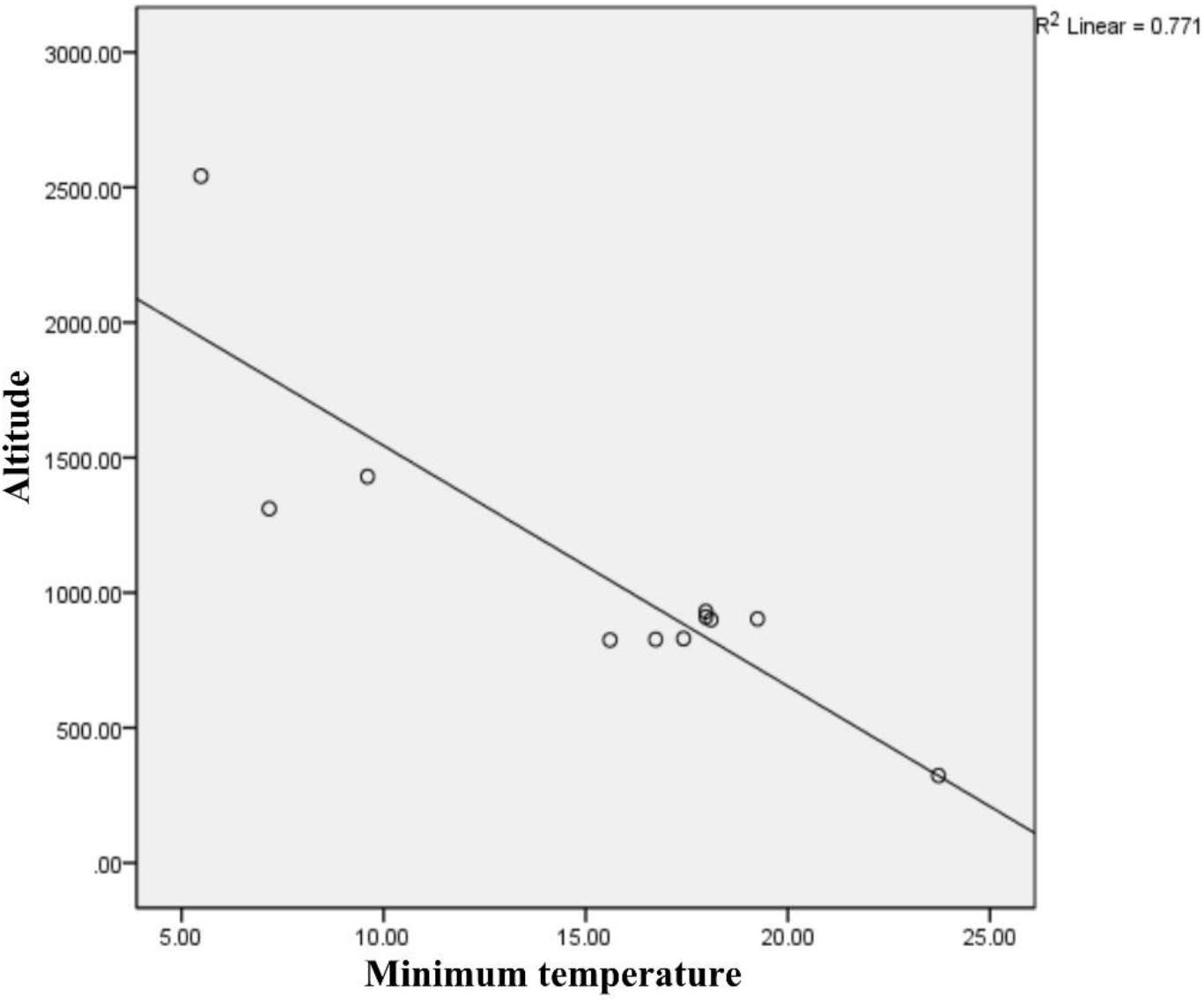
The regression model between altitude and minimum temperature.

**Table 3.** Significant correlation (Carlpearson’s) between different physical parameters from where blood samples of host species were collected.

| Correlations |  |  |  |  |
| --- | --- | --- | --- | --- |
|  |  | Altitude | Minimum Temperature | Relative Humidity |
| Altitude | Pearson Correlation | 1 | -.878** | .347 |
|  | Sig. (2-tailed) |  | .000 | .296 |
|  | N | 11 | 11 | 11 |
| Minimum Temperature | Pearson Correlation | -.878** | 1 | -.282 |
|  | Sig. (2-tailed) | .000 |  | .401 |
|  | N | 11 | 11 | 11 |
| Relative Humidity | Pearson Correlation | .347 | -.282 | 1 |
|  | Sig. (2-tailed) | .296 | .401 |  |
|  | N | 11 | 11 | 11 |

\*\*. Correlation is significant at the 0.01 level (2-tailed).

**Figure 5.**
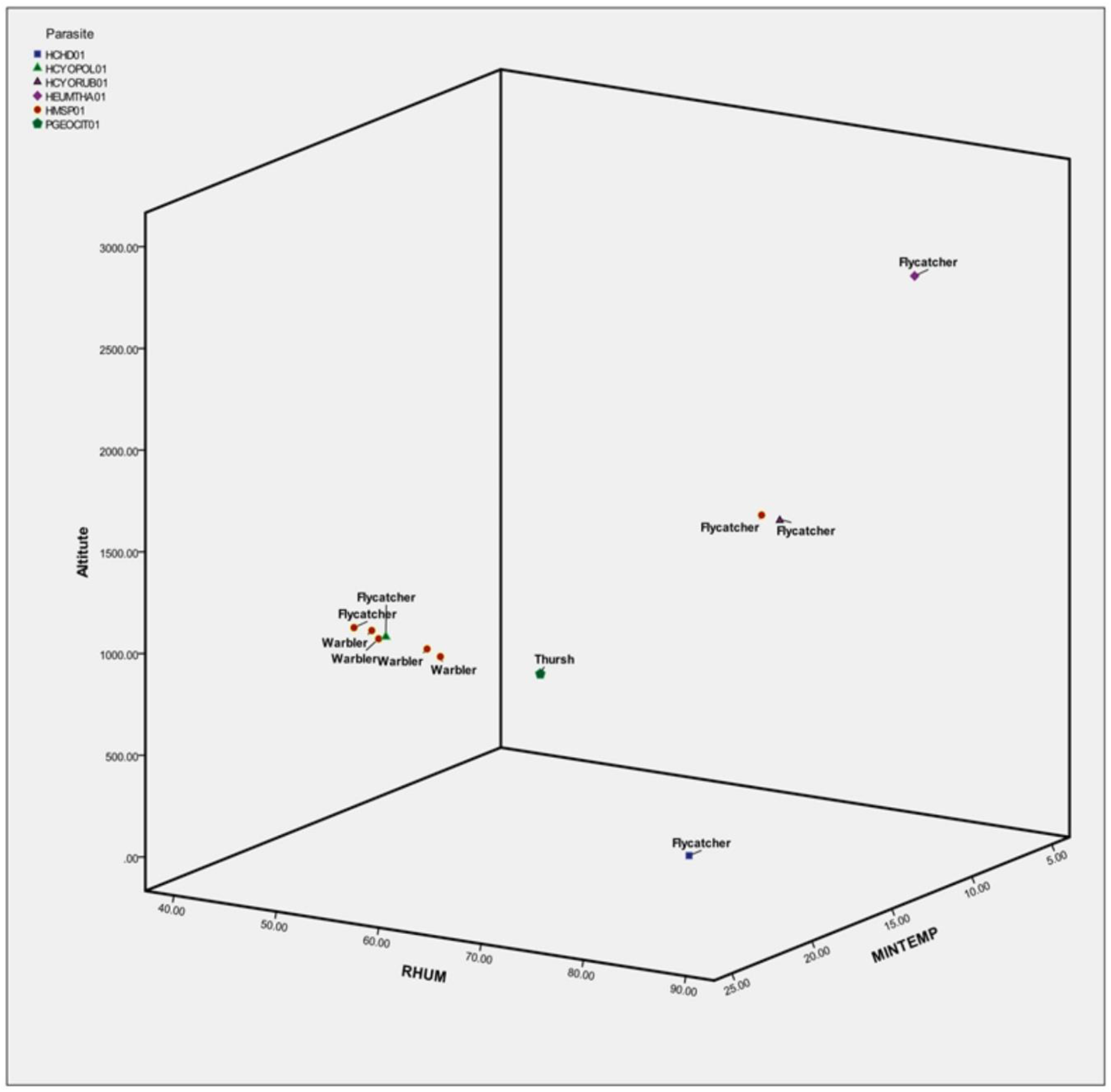
Distribution of host species in relation to altitude, minimum temperature, and relative humidity (RH).

### Spatial phylogenetics

The spatial phylogeny map shows how the haemosporidian lineages of the current study are phylogenetically connected to the other known *Haemoproteus* and *Plasmodium* lineages in India and the world. The role of associated host species in parasite distribution is also discussed. The genetically closest parasite lineage has been shown connected through the coloured tree branch on the globe and the blue coloured polygon area represents the proportional lineage numbers maintained in that geographical region. The lineage MSP01 which was detected in host species *Phylloscopus trochiloides* (LM1, LM2, LM,3, LM4), *Cyornis poliogenys* (LM10) and *Cyornis rubeculoides* (AS89) belonging to the families of Phylloscopidae and Muscicapidae i.e. the groups of Warblers and Flycatchers, respectively (Table 1), are found to be connected through orange tree branches at places between Nagaland and Andhra Pradesh (Fig. 6 A). The *Phylloscopus trochiloides* are strongly migratory, which breeds in northeastern Europe, temperate to subtropical parts of Asia, and winters in India (41). *Cyornis rubeculoides* are native to India, Lao People’s Democratic Republic, Myanmar, Thailand, and Viet Nam (42) and *Cyornis poliogenys* are native to Bangladesh, Bhutan, China, India, Myanmar, and Nepal (43). At the global level, the COLL2 and PFC1 infections were observed in 23 species belonging to the eight families of birds (Muscicapidae, Tyrannidae, Turdidae, Phylloscopidae, Parulidae, Locustellidae, Sylviidae, Ptilonorhynchidae) in the group of Flycatchers, Thrushes, and Warblers and more than 60 percent species out of them were found to be migratory (MalAvi database and present study). So, there are chances, but not sure, that *Cyornis poliogenys* and *Cyornis rubeculoides* might have acquired the *Haemoproteus* infections from species like *Phylloscopus trochiloides* or other which are migratory (more detail in the discussion section) and frequently visiting India from the northern hemisphere. The spatial map showed that most of the *Haemoproteus* lineages of COLL2 and PFC1, present in different hosts mentioned above, were maintained in fourteen countries which were confined in nine polygons (P1-P9) namely India (P1-P3); Malawi (P4); United Kingdom and Netherlands (P5); Finland and Sweden (P6); Russia (P7); Poland, Czechia and Hungary (P8); Alaska (P9), Spain, Malaysia and Philippines, are genetically connected to each other through orange tree branches (Fig. 6B). Figure 6B also represents a complete picture on the genetic relatedness of COLL2 and PFC1 lineages along with their hosts round the globe. The lineages CYOPOL01, EUMTHA01, CYORUB01, and CHD01 have been shown connected to maximum sequence similarity *Haemoproteus* lineages through blue colour tree branches (Fig. 6A). The *Plasmodium* lineage GEOCIT01 relates to PADOM16, through a red branch which was found in *Passer domesticus* from Egypt (Fig. 6C), however, their DNA sequence similarity is incredibly low i.e., 93.89 % (Table S4).

**Figure 6A.**
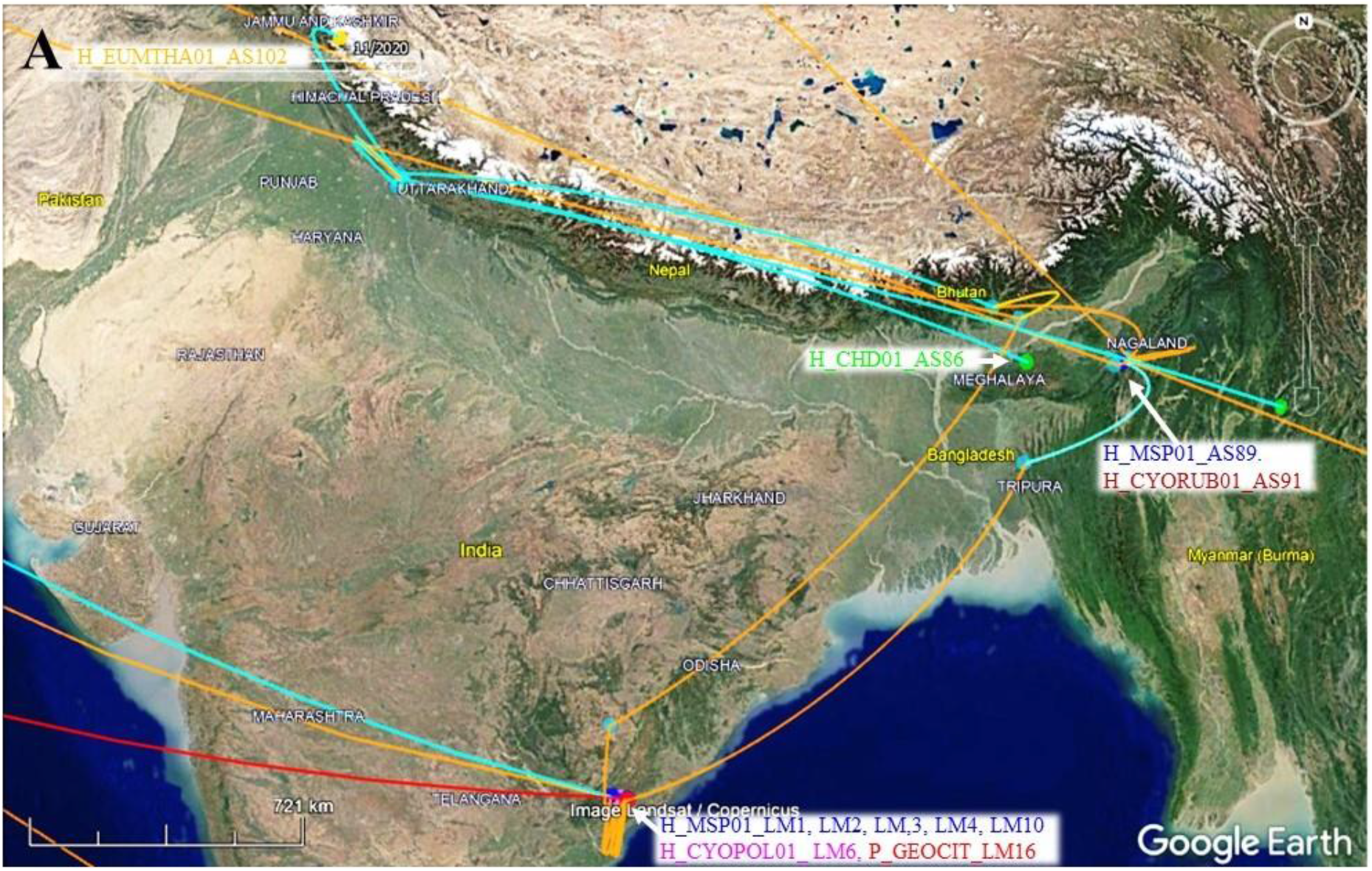
Map showing lineage MSP01 detected in host species *Phylloscopus trochiloides* (LM1, LM2, LM,3, LM4), *Cyornis poliogenys* (LM10) and *Cyornis rubeculoides* (AS89) connected with orange tree branches. Lineages CYOPOL01, EUMTHA01, CYORUB01, and CHD01 are connected to maximum sequence similarity *Haemoproteus* lineages through blue colour tree branches. The lineage on the map and their names have been shown with common colours.

**Figure 6B.**
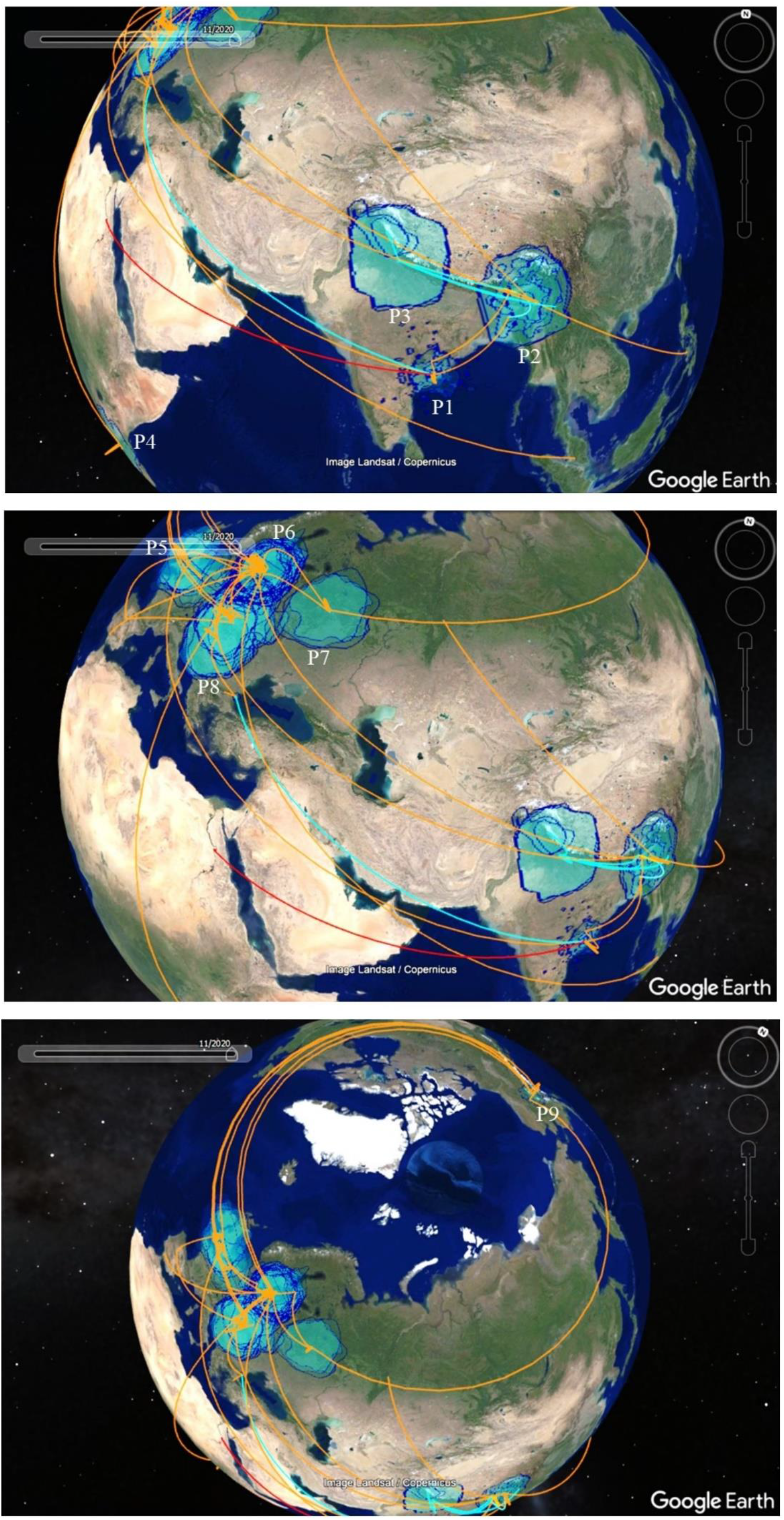
The spatial map shows *Haemoproteus* lineages of COLL2 and PFC1 in nine, blue colour, polygons India (P1-P3); Malawi (P4); United Kingdom and Netherlands (P5); Finland and Sweden (P6); Russia (P7); Poland, Czechia and Hungary (P8); Alaska (P9), Spain, Malaysia and Philippines, connected with each other through orange tree branches. The map also shows a complete picture of genetic relatedness of COLL2 and PFC1 lineages on the globe. Here, ‘P’ stands for polygon.

**Figure 6C.**
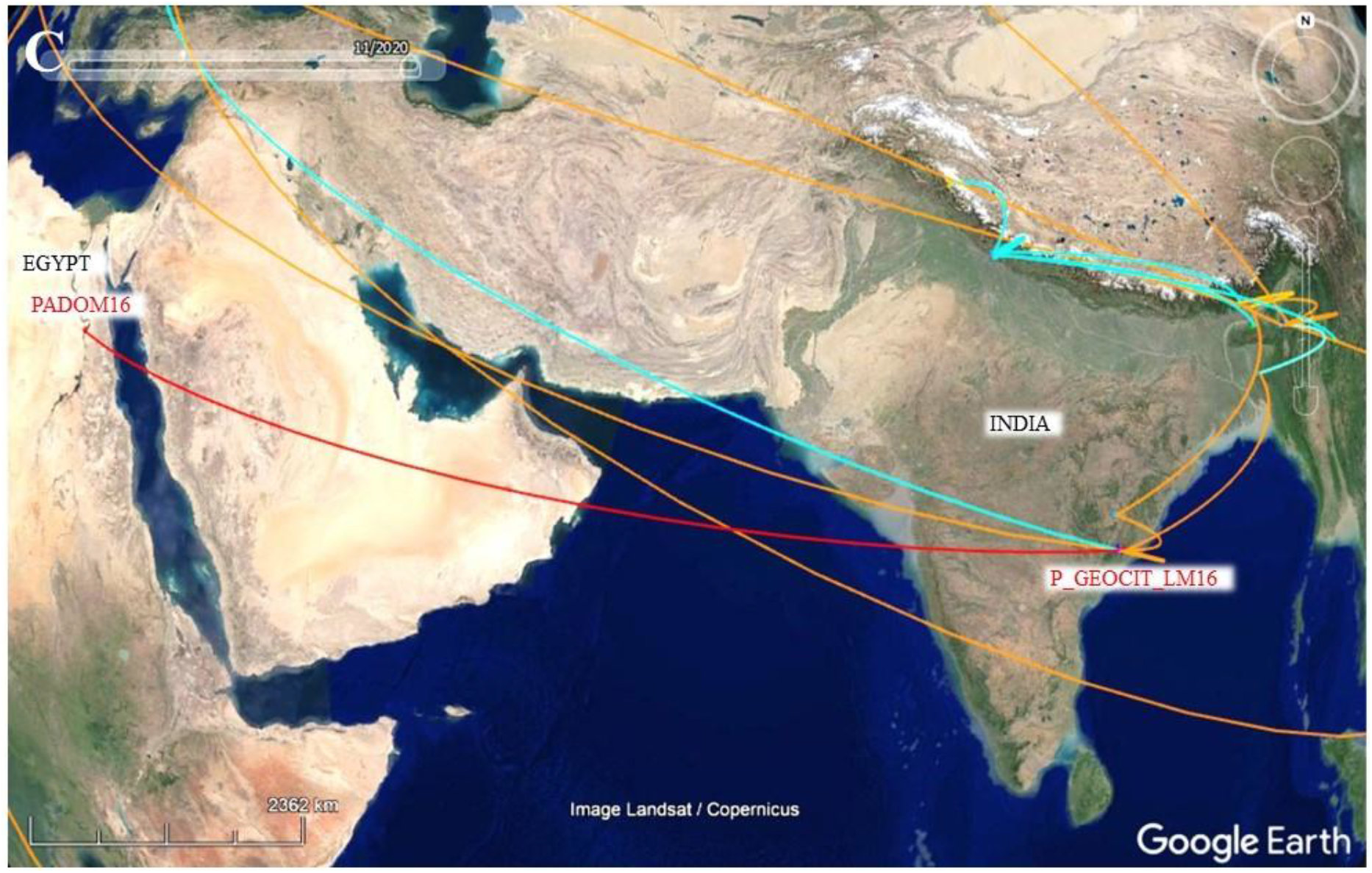
Map shows *Plasmodium* lineage GEOCIT01 in India is linked to PADOM16 (found in *Passer domesticus*) through a red branch in Egypt.

## Discussion

The MSP01 lineage of *Haemoproteus* was detected in *Cyornis poliogenys* (Pale-chinned Flycatcher), *Cyornis rubeculoides* (Blue-throated Blue Flycatcher), and *Phylloscopus trochiloides* (Greenish Warbler) in India. The *C. poliogenys and C. rubeculoides* have been collected from Lanka pakala, Andhra Pradesh and Jotsama, Nagaland, respectively, and both these species are residents of Northeast India (42, 43). *P. trochiloides* is migratory (44) and is found breeding in Afghanistan, Austria, Belarus, Bhutan, China, Czechia, Estonia, Finland, Germany, Kazakhstan, Kyrgyzstan, Latvia, Lithuania, Nepal, Poland, Russian Federation, Slovakia, Sweden, Tajikistan and Ukraine (42). The MSP01 sequence is 100% (386 bp) similar to COLL2 lineage of *Haemoproteus*, so it may be possible that *P. trochiloides* had got infection, initially, from hosts having COLL2 infection during migration. Because it has clearly been shown in the results that COLL2 and PFCI lineages were detected from the hosts present in most of the places (Czechia, Finland, Poland, Russia, and Sweden) where *P. trochiloides* has been reported to migrate (Fig. 6B). The chances of parasites being evolved in-situ or locally are equally likely, however, their maximum sequence similarity with lineages, anywhere in the world, further boosts the above assumption of a source population of getting infection. Other migratory host species with COLL2 infection from Asia, Africa, Europe, and North America vising Indian parts may also be the suspects of spreading MSP01 infection in the above three species. The phylogeography maximum clade credibility tree shows different host species with COLL2 infection in Asia, Africa, Europe and North America connected to MSP01 detected *C. poliogenys, C. rubeculoides* and *P. trochiloides* in Northeast and Andhra Pradesh parts of India through orange colour clades (Fig. 6B).

The comparison of MSP01 cytochrome *b* sequence (387bp) with sequences of morphologically identified *Haemoproteus* species available at the MalAvi database revealed 100% similarity with *Haemoproteus pallidus* (Table S4). PFC1 was also a lineage of *H. pallidus* and MSP01 sequence was also matched with PFCI in the present study. Therefore, hosts having PFC1 infection in different parts of the world may also be the source of *H. pallidus* infection in Indian hosts showing MSP01 infection. However, we could not examine the blood of host species for morphological identification of blood parasites because samples were collected on Whatman filter paper.

The MalAvi database revealed that HCOLL2 lineage is primarily prevalent in 58% of Flycatchers belonging to the two families (Muscicapidae and Tyrannidae), 30% Thrushes family Turdidae, and 12% in Warblers belonging to the four families (Phylloscopidae, Parulidae, Locustellidae and Sylviidae) (Table S5). The PFC1 has been reported from Flycatchers belonging to the family of Muscicapidae (Table S5). The present study results are also in compliance with these as MSP01 is reported in birds belonging to Flycatchers and Warblers (Table S5 and Fig. 5). So, if there is any host specificity factor in these two *Haemoproteus* haplotypes is a topic of future research. It is an interesting fact that the HCOLL2 and PFC1 have only been found associates with Flycatchers, Thrushes, and Warblers in literature, and as the sequence of MSP01 is matching completely with these two lineages, the fact goes same with the current study also. Hence, if there is any host specificity for these three lineages is a matter of future research. The information about the host species with *Haemoproteus* COLL2 and PFC1 infection in different parts of the world has been given in Table S6.

Four unique lineages CYOPOL01, CHD01, CYORUB01, and EUMTHA01 of *Haemoproteus* have been found in *Cyornis poliogenys, Cyornis hainanus dialilaemus, Cyornis rubeculoides*, and *Eumyias thalassinus*, respectively. The DNA sequences of CYOPOL01 is matching more than 99 percent with COLL2 and PFC1; however, it is very difficult to say that it belonged to *Haemoproteus pallidus* until a detailed morphological study is carried out. The same was the case with CYORUB01, of which DNA matched more than 98 percent with COLL2. The other *Haemoproteus* (CHD01 and EUMTHA01) and *Plasmodium* (PADOM16) lineages are connected with distantly related lineages in different parts of the world through other than orange tree branches in the spatial phylogeny map and may not indicate the source, host and parasite populations of infection. The record of EUMTHA01 in *Eumyias thalassinus* is the first of any *Haemoproteus* lineage from the Union Territory (UT) of Jammu and Kashmir. Therefore, a detailed study of blood parasites in Flycatcher and other bird species should be carried out in this UT region. It is very important to discuss here that while making a spatial phylogeny map, we have only used the maximum sequence similarity lineages from the MalAvi database, which had the geographical coordinate information (69.23 percent, Table S6). So, there are equal chances that the infection source may be related to the rest of the hosts and parasite populations and the spatial phylogeny map might be indicating the least sources of infections in host samples of the current study.

The low parasite prevalence seen in the present study might be attributed to multiple factors i.e., PCR failure, host-parasite association, vector distribution, host migration and low sample size. We could only amplify the *Plasmodium* genus-specific fragment of cytochrome b gene (596 bp) using primers L1 and L2 in DNA samples, which also amplified 478 bp fragment using the primers HEAMF and HAEMR2. So, the DNA samples which could not amplify HEAMF and HAEMR2 primers suggested that those were negative for the presence of parasites and there was no PCR failure. The host species’ distribution range was found to be directly associated with the parameters viz. altitude, minimum temperature, and relative humidity (Table 3 and Fig. 5). How the host-parasite association and host migration may affect the parasite prevalence have been explained above in para third and four, respectively. Though we have not collected the parameters responsible for vector distribution in the present study, a lot of information already available in the literature can be utilized for this purpose. The geographic and climatic diversity in India influences the distribution of malaria vectors and parasite species (45). So, malaria endemicity (in the human) is ten times or even more in eastern and southern parts than the extreme northern parts of India (45). No detection of *Plasmodium* from the samples collected from Jammu and Kashmir may be due to the low sample size (n = 11) or distribution status of avian malaria vectors. All culicines of the genera *Anopheles, Culex and Aedes* have been found to act as vectors of avian malaria (46, 47). The records of *Culex* and *Aedes* species were found to be fair enough in comparison to *Anopheles* species in Jammu and Kashmir (48, 49), which suggests low sample size may be the reason for non-detection of *Plasmodium* in this state. The vectors for avian Haemoproteidae belong to Hippoboscidae and Ceratopogonidae families representing hippoboscid flies and biting midges, respectively (20, 2). The records of both the hippoboscid flies (50) and biting midges have been found throughout India (51), may be due to these lineages of *Haemoproteus* were found in all regions of sampling in India.

## Conclusions

In conclusion, the *Haemoproteus* lineage P_MSP01 was found to be more distributed in the samples of this study. Though, the lineage P_MSP01 was genetically closer to the *H. pallidus* generalist lineages H_COLL2 and PFC1 but morphological analysis is required for the final species confirmation. The *H. pallidus* is known to have a wide geographic distribution and host species range (Ortiz-Catedral et al., 2019) and our results are also supporting this. The genetic relatedness of *Haemoproteus* lineages P_MSP01 in India with COLL2 and PFC1 (supported by high posterior probability values) suggest possible infection, through migratory host species, from Africa, Europe, North America, Malaysia, and Philippines. The physical factors (altitude, minimum temperature, and relative humidity) were associated with the host species’ distribution range.

## Supporting information

Table S1. The details obout the bird samples collected from India.

Table S2A All lineages of Plasmodium (n=1287) and Haemoproteus (1575) downloaded from MalAvi database for avian haemosporidians parasites, as on 24-05

Table S2B Total lineages of Plasmodium (1230), Haemoproteus (1451) and Leucocytozoon (03) whose cytochrome b gene (230 bp) sequences were finally used

Table S3. Maximum percent similarity lineages of Plasmodium and Haemoproteus (only having GPS coordinates), obtained from MalAvi data base, which were

Table S4. Maximum percent cytochrome b gene sequence similarity of lineages from current study with lineages from MalAvi data base for avian haemospor

Table S5. Prevalence of COLL2 and PFC1 in host species at MalAvi database and present study.

Table S6. All Plasmodium and Haemoproteus lineages, having maximum percent similarity with lineages from current study, downloaded from MalAvi data ba

## Acknowledgments

The current study was part of the project under UGC-Dr. D.S. Kothari Postdoctoral Fellowship Scheme, UGC awarded to the first author (BL/16-17/0151) from March 2017 to March 2020. We are thankful to National Institute of Malaria Research, Dwarka, New Delhi, for sparing the positive *Plasmodium* DNA sample for the standardization work. We acknowledge the support of the Director, Indian Institute of Integrative Medicine, CSIR, Jammu and Dr. Sumit Gandhi, Principal Scientist, for helping us to carry out the standardization work of this project. We sincerely thank Professor Trevor Price, University of Chicago for his generous support. We thank Forest department of Union Territory of Jammu & Kashmir, Himachal Pradesh, Uttarakhand, Sikkim, Arunachal Pradesh, Mizoram, Andhra Pradesh and Meghalaya for granting permission for sample collection.

## Authors contributions

Conceived the idea and designed the experiments: V NKT AS. Performed the experiments: V AS BS. Analysed the data: V AS VS. Contributed reagents/ materials/ analysis tools: SKG V AS RD DM. Wrote the paper: V VS SKG. Reviewed and edited the paper: SKG NKT DM CPS RD.

## Ethics statement

The bird samples, whose DNAs were used in the present study, were collected with due permissions of the respective state forest departments.

## Data availability

All data related to this paper has been submitted to this journal and is available to the readers.

## Notes

### Competing Interest Statement

The authors have declared no competing interest.

### Summary of Updates

Format of the manuscript is improved.

