## Supplementary material for "Avian haemosporidians (*Plasmodium* and *Haemoproteus*) status in selected bird groups (Old world Flycatchers, Warblers, Babblers, and Thrushes) of India and their phylogenetic relationships with other lineages of the world": Table S4. Maximum percent cytochrome b gene sequence similarity of lineages from current study with lineages from MalAvi data base for avian haemospor

Table S4. Maximum percent cytochrome *b* gene sequence similarity of lineages from current study with lineages from MalAvi data base for avian haemosporidians parasites.

| Lineage from the current study | Maximum DNA sequence % similarity with lineages from MalAvi data base | | Maximum DNA sequence % similarity with morphologically identified species on MalAvi data base | | | |
| --- | --- | --- | --- | --- | --- | --- |
|  | 230 bp | | 230 bp | | 384 bp | |
| MSP01 | H_COLL2 (H_Pallidus) | 100 | H_COLL2 (H_Pallidus) | 100 | H_COLL2 (H_Pallidus) | 100 |
|  | H_PFC1 (H_Pallidus) | 100 | H_PFC1 (H_Pallidus) | 100 | H_TUCHR01 (H_Minutus) | 99.75 |
|  | H_NEOPOE02 | 100 |  |  |  |  |
|  | H_SYCUR01 | 100 |  |  |  |  |
|  | H_SYCUR02 | 100 |  |  |  |  |
|  | H_TUPHI01 (H_Minutus) | 100 | H_TUPHI01 (H_Minutus) | 100 |  |  |
|  | H_TUROLI05 | 100 |  |  |  |  |
| CYOPOL01 | H_COLL2 | 99.12 | H_COLL2 (H_Pallidus) | 99.56 |  |  |
|  | H_PFC1 (H_Pallidus) | 99.56 |  |  |  |  |
| CHD01 | H_TROERY02 (H_Homoleithricus) | 97.38 | H_COLL2 (H_Pallidus) | 96.51 | 10935 |  |
|  | H_TURSTR03 | 97.38 |  |  | 11368 |  |
| CYORUB01 | H_COLL2 | 98.69 | H_COLL2 (H_Pallidus) | 98.69 |  |  |
| EUMTHA01 | H_CARCHL06 | 99.13 | H_COLL2 (H_Pallidus) | 98.69 |  |  |
| P_GEOCIT01 | P_PADOM16 | 93.89 | P_PADOM16 | 93.89 |  |  |
